# Local trade-offs shape flower size evolution across Arabidopsis thaliana distribution

**DOI:** 10.1101/2025.04.19.649637

**Authors:** Kevin Sartori, Clàudia F. Mestre, Md Jonaid Hossain, Aurélien Estarague, Elza Gaignon, Pierre Moulin, Jesse R. Lasky, Denis Vile, Francois Vasseur, Cyrille Violle, Adrien Sicard

## Abstract

The diversification of flowers, largely driven by mutualistic interactions with animal pollinators, has generated remarkable variation in floral form and size and is thought to have driven the evolutionary radiation of angiosperms. Here, we investigate the geographic variation in reproductive organ growth in the self-fertilizing species *Arabidopsis thaliana*, where the loss of pollinator dependence is predicted to favour reduced floral investment through resource reallocation. We find extensive variation in flower size underpinned by a polygenic architecture, in which derived alleles both increase and decrease petal size and show signatures of positive selection. Nevertheless, the evolutionary direction of flower size differs across regions, reflecting environmentally mediated shifts in the relationship between flower size and seed production. Strong purifying selection at the climatic margins favours smaller flowers, whereas relaxed environmental constraints in more suitable habitats allow the emergence and persistence of both small and large-flower variants. This biogeographic pattern extends to other life-history traits, highlighting how environmental heterogeneity and resource allocation constraints shape evolutionary trajectories and maintain phenotypic diversity in selfing lineages.

## Introduction

Natural selection balances flower organ construction and maintenance costs against the likelihood of fertilization (Roddy et al., 2020). These costs follow classical allocation theory, as illustrated by the trade-off between flower size and number (Caruso, 2006; Roddy et al., 2020; Worley & Barrett, 2000). In animal-pollinated outcrossing species, pollinator-mediated selection often favours large and more conspicuous flowers adapted to pollinator nutritional needs and behaviours, promoting flower diversification (Crane et al., 1994; Friis et al., 1994, 1999, 2006; Galen, 1999). Following the transitions to self-fertilization, however, selection is expected to reduce investment in floral displays and reallocate resources to other fitness components (Stebbins, 1957). This mechanism is frequently invoked to explain the characteristic reduction in flower size and attractiveness, collectively known as the “selfing syndrome”, in selfing species (Sicard et al., 2011). Yet, empirical studies quantifying the relative importance of resource-allocation trade-offs compared to other evolutionary processes remain limited.

Different evolutionary processes could influence floral evolution in selfing species, each leading to distinct patterns of variation in flower size. When the construction of floral organs incurs high energetic costs, selection should favour mutations that reduce floral display size, thereby enhancing seed production through resource reallocation (Charlesworth & Charlesworth, 1981). This directional selection scenario predicts (i) a negative association between flower size and seed production, (ii) genomic evidence of positive selection on alleles that reduce flower size, and (iii) a general convergence toward smaller flowers with reduced within-species variance (Stephan, 2019). Because environmental heterogeneity influences both resource availability and selection strength (Troth et al., 2018), these patterns should be more pronounced in resource-limited environments (Fu et al., 2010; MacTavish & Anderson, 2020; Tuller et al., 2018). Alternatively, the loss of attractive traits could confer little to no fitness benefit, and selection maintaining large displays would be relaxed. Under this scenario, the trajectory of floral-size evolution may become less predictable. In this case, the role of genetic drift should play a more dominant role, leading to idiosyncratic shifts in mean size, widening phenotypic variance, and few clear signatures of directional selection at causal loci (Woźniak & Sicard, 2018). Reduced effective recombination under high homozygosity can further facilitate drift-mediated fixation of nonsynonymous mutations that impair floral-growth genes (Barrett et al., 2014).

Additional processes may interact with these trajectories. Pleiotropy can couple floral size variation with other organs controlled by shared developmental pathways, producing correlated responses that range from near isometric to clearly allometric, depending on the extent of shared genetic control (Huxley et al., 1993; Johnson & Lenhard, 2011). Likewise, linked selection, enhanced in selfers due to low effective recombination, can further reshape diversity around selected loci and indirectly hitchhike alleles affecting floral size (Slotte, 2014). Given the diversity and nature of the mechanisms involved, floral-size variation is likely to be highly polygenic, consistent with observations in selfing lineages (Abraham et al., 2013; Sicard et al., 2011; Sicard & Lenhard, 2011; Woźniak et al., 2020). However, the relative contributions of these mechanisms remain poorly understood.

*Arabidopsis thaliana* provides a powerful system to test these predictions. The evolution of self-fertilization in this species occurred relatively recently and was followed by rapid range expansion across diverse climates and environments (Bechsgaard et al., 2006; Shimizu & Tsuchimatsu, 2015; Tang et al., 2007). This history provides opportunities for environment-specific selection on floral investment and the fixation of distinct size-affecting mutations across lineages (Tsuchimatsu et al., 2020). Leveraging extensive genomic and phenotypic resources, we examine the genetic and ecological drivers of floral-size evolution following the emergence of selfing and assess how environmental gradients shape these evolutionary trajectories across the species’ range. Our results reveal that patterns of phenotypic evolution differ geographically, reflecting variation in selection pressures across environments.

## Materials and Methods

### Biological materials

A subset of 407 *Arabidopsis thaliana* accessions was selected, overlapping the complete range of genetic structure and geographic distribution represented in the 1001 genome project accessions (Alonso-Blanco et al., 2016) (**Fig. S1, Dataset S1**). All accessions were vernalized for 4 weeks in the dark at 5°C in October 2022. Seedlings were then transferred to soil in individual pots, with one plant grown per accession, and grown in a glasshouse under semi-controlled conditions in Montpellier, CEFE CNRS, France (maintained at around 20°C during the day and 15°C at night with natural light). Accessions were randomly distributed on three tables and daily rotated to homogenize the effect of spatial heterogeneity within the glasshouse across the genotypes. Plants were surveyed daily, and the date of first flower opening, flower collection and plant death were recorded. The fourth rosette leaf was collected at bolting and immediately scanned. Three flowers were collected between the 10th and the 25th flower of the main stem from the primary inflorescence. Flowers were collected and preserved in 70% ethanol in individual Eppendorf tubes placed in the dark at room temperature before measurements.

### Morphological measurements

Flowers were stained with 0.02% Toluidine Blue for 10 minutes before being dissected and placed in between microscope slides and cover slips with a drop of glycerol (40% v/v). Slides were scanned at a 4800-dpi resolution with an Epson perfection v850 pro® scanner. Leaf, petal, sepal, and short and long stamen sizes (length, width, and area when relevant) were measured using Fiji version 2.35 (Schindelin et al., 2012). The blue channel of the images was isolated, and contrast-enhanced with Otsu’s local contrast method. Leaves were scanned at 800-dpi resolution and made binary. Pixels that belong to the organs were counted to estimate organ sizes. Measures were calibrated using the dots per inch (dpi) of the images. The ovule number per flower was counted manually using fluorescence microscopy (Leica DMI4000). Flowering time was calculated as the duration in days from seed sowing to the opening of the first flower. We computed an adjusted trait values by extracting the residuals from a linear model that included the relevant confounding factors (petal rank along the stem, stem number, and a variable describing spatial distribution in the tables). These adjusted values were used for all downstream statistical analyses, including GWAS. All traits were normally distributed and did not necessitate transformation (**Figure S2**).

### Data collection from the literature

Taking advantage of the extensive literature available about the model species, we confronted our phenotype data to published datasets that shared a significant number of accessions. In particular, we used the 107 phenotypes gathered by (Atwell et al., 2010), containing phenology, microbial resistance, plant morphology and ions content values, and the Aradiv database from (Przybylska et al., 2023) that contains plant functional strategies related traits. We also used datasets reporting fitness in terms of seed production, where a majority of our accession set was shared (Exposito-Alonso et al., 2018; Wilczek et al., 2014), and compared our measurement of flower organ size with other available datasets (Li et al., 2020; Wiszniewski et al., 2022). In addition, genome-wide nucleotide polymorphism data from the sampled accessions were obtained from publicly available variant calling format (VCF) data from the 1001 genomes project webpage (https://1001genomes.org/data/GMI-MPI/releases/v3.1/) (Alonso-Blanco et al., 2016).

### Statistical analyses

All statistical analyses were performed with R (version 4.3.2). We performed principal component analyses with the package *FactoMineR*. Testing for isometry versus allometry was achieved with standard major axis regression, which assumes an error component from both tested variables, with the package *smatr* (Warton et al., 2006). Skewness and deviation from the average were statistically tested in R using a skewness test and a Student’s t-test, respectively. Triangular relationships were tested by comparing statistically the slopes of 5% and 95% quantile regressions (Peng, 2017) using the R package *quantreg*.

### Genome-Wide Association Studies (GWAS)

We tested associations between organ size traits and genome-wide nucleotide polymorphisms using the GEMMA software v0.98.1 (Zhou & Stephens, 2012). Only single-nucleotide polymorphisms (SNPs) were retained in the association analyses, short indels and non-ACTGN were filtered out, as well as SNPs with a minor allele frequency below 5% using PLINK (Purcell et al., 2007). A total of 1,899,962 SNPs passed the filters. GEMMA takes the population structure into account by computing and including a relatedness matrix as covariate in the models. The P-values distribution of all GWAS followed a uniform distribution and did not necessitate transformation for false discovery rate, and genomic inflation factor tended to 1±0.1.

The *bslmm* function (Bayesian sparse linear mixed model) was used to estimate the proportion of phenotypic variance explained by the genotypes (PVE, the SNP-based heritability of the traits) and the Single Nucleotide Polymorphisms effect sizes, as recommended by Zhou et al. (2013) by the formula α + β.γ (with α the small additive effect of each SNP, β the additional effect of some large effect SNPs and γ its posterior inclusion probability estimate, all returned by the software).

The *lmm* function (linear mixed model) was used to identify specific quantitative trait loci (QTL) associated with the traits. Despite strong PVE detected notably for petal and sepal size traits (PVE > 0.7; Table S2), associations rarely exceeded the arbitrary Bonferroni significance threshold accounting for multiple testing. Under the assumption that this is indicative of the interplay of multiple small-effect QTLs, the threshold of significance was relaxed to consider the potential polygenic architecture of the studied traits. Reports of P-values detecting relevant genes range from 10-7 to 10-4 (Burton et al., 2007; Chen et al., 2021), but there is no consensus on a threshold to detect multiple minor effect variants in a polygenic architecture (also known as multiple disease phenotype). Because the Bonferroni correction can be overly conservative for traits with a polygenic architecture, we adopted an empirical threshold based on the recovery of the well-established flowering-time regulator *FLOWERING LOCUS C* (*FLC*) in the flowering-time GWAS. The threshold corresponding to this association (P = 5.2 × 10⁻⁵) was subsequently applied to identify candidate loci for other traits. FLC, known as a key regulator of flowering time, is commonly detected in GWAS in *A. thaliana*, and its variation accounts for more than half the trait’s variation (Sasaki et al., 2018). To eliminate genetically linked variants, we filtered out significant SNPs by considering their genomic position (https://www.arabidopsis.org), statistical significance, and predicted function using SnpEff (Cingolani, 2022). This process resulted in retaining one focal polymorphism per QTL for subsequent analyses (**Table S**4).

### Population Genetics Analysis

We first assessed the regime of selection acting on leaf and flower size QTLs across the species distribution with two metrics. We first computed the πN/πS ratio for all *A. thaliana* annotated genes. We calculated the ratio between the genes’ nucleotide diversity in non-synonymous sites (πN – defined as the 0-fold degenerate sites) and the nucleotide diversity in synonymous sites (πS – defined as 4-fold degenerate sites). The rationale is that πS gives a measure of neutral genetic variation and serves as a reference for πN, which measures the amount of genetic variation at sites under natural selection control. Thus, a value of πN/πS below 1 is indicative of purifying selection, and a value above 1 is indicative of diverging selection. We used the genome-wide distribution as a reference: individual genes’ πN/πS included in the 0.05 to 0.95 quantile do not deviate from the general purifying trend. Student-test and skew-test were performed on R to evaluate a set of genes πN/πS deviation from the average and normality. Full sequences of gene coding regions and general feature format files were downloaded from Phytozome 13 (version Athaliana_447_TAIR10). Together with the VCF file, they were used to generate the diversity of gene sequences observed among *A. thaliana* genotypes. The R package *RGenetics* was used to discriminate 4-fold from 0-fold degenerate sites, and the *nuc.div* function from *pegas* was used to compute the nucleotide diversity. More details about the methods and scripts used are available at (https://github.com/kevinfrsartori/Whole_Genome_PiNPiS). Second, the extended haplotype homozygosity (EHH) was computed genome-wide to test whether a particular derived variant is subjected to ongoing or partial sweep in populations (Sabeti et al., 2002). When advantageous mutations are selected, the frequency of genetically linked variants also increases, extending the haplotype length around the selected site. This signal perdures until recombination events erase it. EHH measure the decay of a haplotype length upstream and downstream of the focal SNP, which thus depends on the selection’s history and strength. Since the patterns of EHH might also be affected by the species’ population structure and demographic history (Voight et al., 2006), we compute this estimate genome-wide to assess significance. The R package *rehh* (Gautier & Vitalis, 2012) was used to compute the EHH’s test statistic (iHS), defined as the log ratio of the integral of the EHH decay (iHH) around the derived allele divided by the iHH around the ancestral allele. This statistic was then compared to the distribution of iHS for all SNPs of similar allele frequency, and a P-value was reported. The iHS computation relies on the polarization of the SNP dataset, i.e. the ancestral state of each SNP. To differentiate between the derived versus ancestral allele, we used a maximum likelihood method using the focal species allele frequencies together with related species allelic information. We first identified an ortholog gene in *Arabidopsis lyrata* and *Arabidopsis halleri* for most *Arabidopsis thaliana* genes by using OrthoFinder (Emms & Kelly, 2015). We kept all *A. thaliana* genes having at least one ortholog and kept only one ortholog per outgroup species based on the genes’ distance tree, resulting in a list of 25411 genes. We extracted a sequence covering the coding region plus the cis- and trans-regulatory regions for all trio of genes, and performed a multiple alignment using MAFFT (Katoh et al., 2002). We extracted the allelic information of the outgroup species with custom R scripts and ran the program est-sfs (Keightley & Jackson, 2018). It is important to note that the computation of iHS was limited to the presence of ortholog genes, resulting in filtering out few SNP per trait lists (**Table S4**).

### Habitat suitability

The *A. thaliana* niche was modelled using the MaxEnt software (Phillips et al., 2006) by reproducing the method from (Yim et al., 2022). Climatic data were downloaded from CHELSA v2.1 (https://chelsa-climate.org) at a resolution of 30 arc seconds (Karger et al., 2017). The species distribution modelling was performed with the following variable panel: the isothermality (ratio of diurnal variation to annual variation in temperatures, °C), the mean of daily minimum air temperature of the coldest month (°C), the annual range of air temperature (°C), mean daily mean air temperatures of the wettest quarter (°C), mean daily mean air temperatures of the warmest quarter (°C), precipitation seasonality (standard deviation of the monthly precipitation estimates, kg m-2), mean monthly precipitation amount of the wettest quarter (kg m-2 month-1), and mean monthly precipitation amount of the driest quarter (kg m-2 month-1). Altitude data at a resolution of 30 arc seconds come from the Global Multi-resolution Terrain Elevation Data 2010 (GMTED2010) (Danielson & Gesch, 2011) and were downloaded from the USGS website. We used the extensive species occurrence data (N=672) provided by (Yim et al., 2022), which covers the distribution of the species in an evenly spaced manner (one sample per km2) to account for sampling bias. From the MaxEnt model, we estimated the habitat suitability (HS) and the limiting factor (LF) of all *A. thaliana* accessions’ collecting sites (**Figures S6; S7**). The HS metric scales from zero to one, while LF refers to the climatic variable that has the most significant impact on HS when changed while maintaining all others variable constant. The relationships between traits and HS were tested using linear regressions. Linear regressions were plotted when significant, and quantile regressions were plotted when the test for slope difference (Anova) was significant. In order to test for selection regime shift along habitat suitability, we divided the *A. thaliana* accession set into ten equally sized sub-populations. To avoid the influence of over-sampling in some regions of the species distribution, we first pruned the accession list based on genetic relatedness. We used the PLINK software to estimate the number of accessions passing a filtering threshold of genetic relatedness ranging from r=0.1 to r tends to 1. We found that the number of related accessions stops dropping after 0.95 and remains constant while r tends to 1 (**Figure S9**). We thus excluded 87 highly correlated accessions. HS ranges were obtained by dividing the HS distribution of unrelated accessions into ten quantiles, where each quantile contains about 100 accessions.

### Selection regime along habitat suitability ranges

The HS range samples obtained by splitting the *A. thaliana* accessions into 10 populations did not have a geographic and historic meaning for the species, which prevents from computing most classical population genetic metrics. Thus, we tested for changes in selection pressure by relying on allele frequency only. We used the sequencing dataset presented above and relaxed the minimum allele frequency threshold to a minimum allele count of 1. Only the SNPs having ancestry information located in the gene body were kept, and we computed the unfolded Site Frequency Spectrum (uSFS) for leaf and flower organs associated genes. The allele counts for ranges of allele frequency of 0.05 were reported. Since new mutations are putatively deleterious, natural selection maintains new allele frequencies at a low level. As a result, uSFS are right-skewed distributions of allele counts, with a large number of low-frequency mutations. The skewness of the distribution can be reduced by a drop in purifying selection.

## Results

### Unsuspected variation of flower size in A. thaliana is highly heritable

The directional selection hypothesis predicts that both the mean and variance of flower size decline in self-fertilizing species such as *Arabidopsis thaliana* due to resource allocation trade-offs that favour reduced investment in floral display. Alternatively, genetic and phenotypic variation may persist if drift predominates (Buysse et al., 2024) or if spatially variable environments maintain divergent selection (MacTavish & Anderson, 2020).

To test these predictions, we measured the length, width, and area of floral organs and leaves in 407 *A. thaliana* accessions spanning the species’ global genetic and geographic diversity (**Fig. S1**; **Dataset S1**). Variance-based heritability estimates (**Dataset S2**), accounting for trait variance both within and between regional populations of *Arabidopsis thaliana* (Alonso-Blanco et al., 2016), revealed high heritability for all investigated traits. In particular, petal and sepal traits showed very high values (H² > 0.8), indicating that most of the observed variation is genetically determined and only minimally influenced by environmental factors. All traits were variable and approximately normally distributed (**Fig. S2**). Floral organ size explained most phenotypic variance among accessions (43.2%, PC1; **Fig. 1a**), with strong correlations among floral traits (r > 0.5). Leaf size variation dominated PC2 (21.6% of the variance). Ovule number contributed mainly to PC4 (<8% of variance). Petal area showed the greatest variability, matching leaf area in its coefficient of variation and spanning more than a threefold range (**Dataset S2**). Moreover, we compared our petal size measurements with independent flower diameter estimates. We detected significant correlations, with genotype rankings from small to large flowers remaining consistent across treatments. (Wiszniewski et al., 2022) and across experiments (Spearman’s rank correlation coefficient ≈ 0.46), indicating that relative differences among genotypes are largely preserved across environments

**Figure 1:**
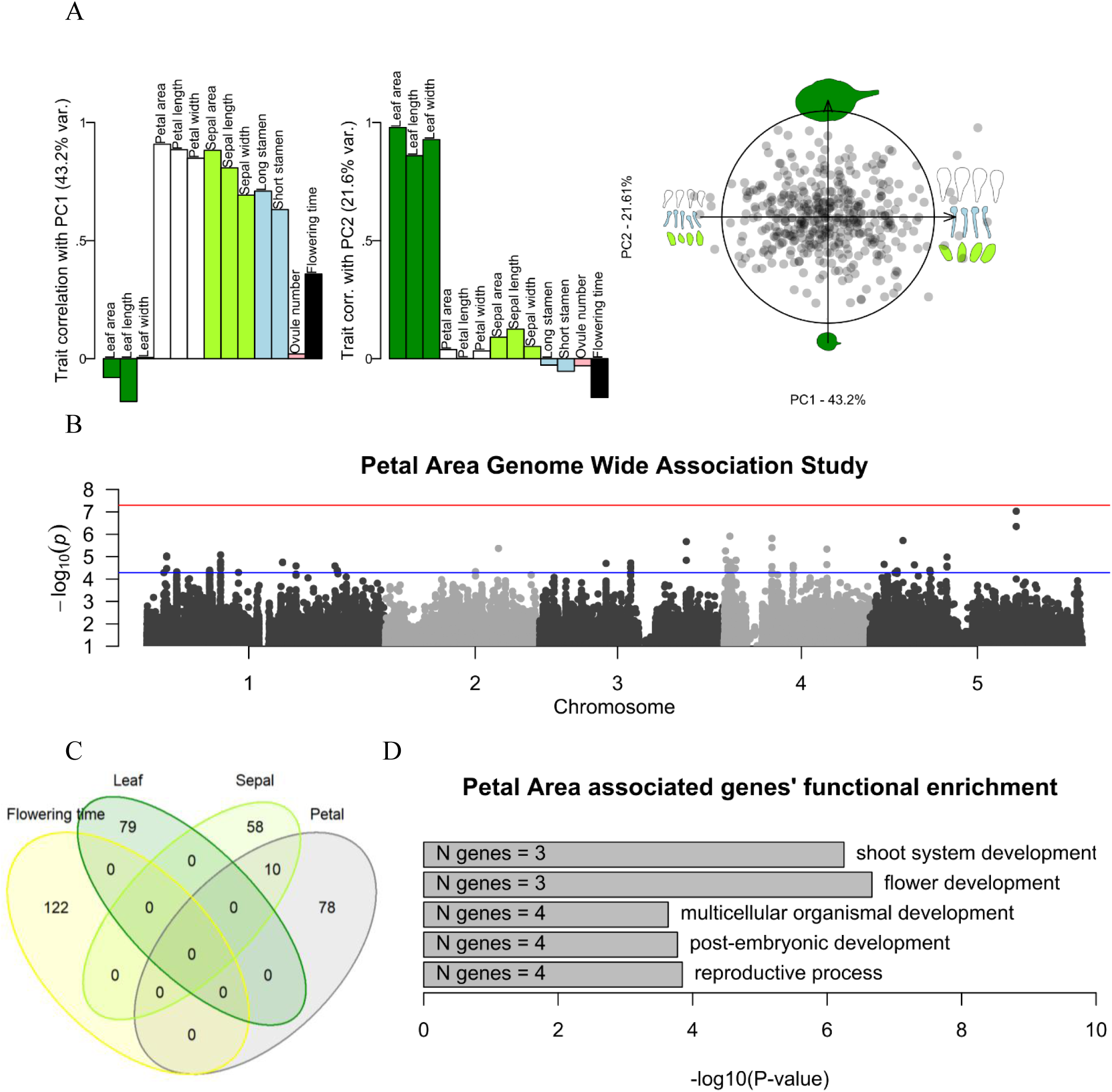
Extensive flower size variation in *A. thaliana* is determined by a polygenic architecture of low pleiotropic mutations. **A)** A Principal component analysis (PCA) was performed on flower organs and leaf size excluding flowering time that was correlated post hoc and represented as a supplementary variable. Bar plots show trait correlations with principal components (PC), with colours indicating different organs: dark green, leaf; light green, sepal; white, petal; blue, stamen; pink, ovules. The biplot displays accession coordinates along the first two principal components (PC), with organ silhouettes representing extreme organ size values at the same scale; **B)** Manhattan plot of the petal area genome-wide association study. Red line represents the Bonferroni threshold, and the blue line represents the relaxed polygenic threshold. **C)** Venn diagram illustrating the lack of overlap between genes associated with leaf and flower organs’ size and flowering time variation. D) Representation of the gene ontology functional enrichment of petal area associated genes. Only significant terms are represented, including gene number and P-value of a fisher exact test.

These results indicate that floral-size variation in *A. thaliana* remains substantial despite selfing, consistent with either incomplete fixation of size-reducing alleles during the evolution of selfing or environmentally structured selection maintaining divergence among lineages. Combined with the high heritability estimates, this suggests that the observed phenotypic differences largely reflect stable genetic differences among accessions, supporting the use of this dataset to dissect the genetic basis of natural variation in petal size.

### Independent, low-pleiotropy mutations largely account for the variation in flower size

We investigated whether indirect effects, such as linked selection (Smith & Haigh, 1974) or pleiotropy with other traits under selection, drive the evolution of floral size in *Arabidopsis thaliana*. To this end, we first tested for phenotypic covariation between floral organ size and other traits among genotypes (**Table S2**). Stamen, petal, and sepal size were significantly and positively correlated (r = 0.28–0.93) but showed no correlation with ovule number or leaf size, except for a weak negative correlation between leaf length and petal width (r = –0.15; **Figure S3**). The relationship between long-stamen length and petal length was strictly isometric (slope = 1, P-value > 0.01), suggesting an overlap in their genetic bases, while other pairwise relationships were allometric (slope < 1, P-value < 0.001). All floral organs were positively correlated with flowering time, although the correlation coefficient did not exceed 0.5 (r = 0.14–0.42). We further examined correlations with published phenotypes. Floral organ size traits showed no significant correlation with any of the 107 phenotypes from Atwell et al. (2010) (**Table S2**). Specific leaf area (SLA), leaf nitrogen content (LNC), and total plant biomass (Przybylska et al., 2023) were significantly correlated with several floral size traits from our dataset (r = –0.35–0.4). Most traits from (Przybylska et al., 2023) were also correlated with our measurement of flowering time (r = –0.67–0.70). To determine whether phenotypic correlations result from shared genetic bases, we quantified genetic correlations using multivariate GWAS (mGWAS) for significantly correlated trait pairs. The mGWAS partitions genetic variance into trait-specific and shared components, allowing estimation of the genetic correlation (r_g_) between traits (Zhou & Stephens, 2014) (**Table S3**). Within our dataset, we detected no significant genetic correlations between floral and leaf traits, indicating independent genetic bases. Petal and sepal areas showed weak but significant correlations with LNC and plant dry mass (rg ∼ 0.3, P < 0.01), which are proxies for resource acquisition ability (Pérez-Harguindeguy et al., 2013). These results suggest that resource acquisition capacity modestly influences both vegetative carbon economy and floral organ growth, supporting a role for allocation constraints in flower-size evolution. Overall, floral organ size variation showed limited phenotypic and genetic correlation with other traits, indicating that pleiotropy and linked selection exert only minor influence on floral size evolution in *A. thaliana*.

We next conducted univariate GWAS to identify loci underlying individual trait variation (**Fig. 1b**). We identified 61, 105, 87, 59, 38, and 65 SNPs, corresponding to 144, 191, 194, 126, 117 and 122 candidate genes, associated with leaf, petal, sepal, stamen size, ovule number and flowering time respectively (**Fig. 1c**; **Dataset S4**). Functional annotation using TAIR gene ontology terms (Huala et al., 2001) identified several known growth regulators among candidate genes, including *JAGGED*, *GROWTH-REGULATING FACTOR 6*, *CYTOCHROME P450 90B1*, *KINASE-INDUCIBLE DOMAIN INTERACTING 9*, *E2FE*, *MYB DOMAIN PROTEIN 104*, *RESPONSE REGULATOR* 7, *INDOLE-3-ACETIC ACID INDUCIBLE 19*, and *SEEDSTICK* (Dubos et al., 2010; Gómez-Ocampo et al., 2023; Gonzalez et al., 2015; Lee et al., 2008; Lopes et al., 2023; Maki et al., 2022; Omidbakhshfard et al., 2015; Sauret-Güeto et al., 2013; Vlieghe et al., 2005). Genes associated with petal area were significantly enriched for functions related to floral and shoot development (6 genes, P ∼0.01) and reproductive regulation (12 genes, P ∼ 0.02; **Fig. 1d**), thereby supporting our ability to detect the polygenic architecture of the trait. We observed no overlap between SNPs and genes associated with floral and leaf area, consistent with the absence of phenotypic and genetic correlations (**Fig. 1c**). However, we detected among petal area associated genes, the MADS-box transcription factor FLOWERING LOCUS M (FLM), first identified as a regulator of flowering and later associated to more complex pleiotropic effect on plant organ growth (Hanemian et al., 2020). Despite moderate phenotypic correlations among floral organs, their genetic determinants showed limited overlap. The observed phenotypic covariation is likely reflecting coordinated responses to shared environmental conditions rather than shared genetic architecture. Thus, floral organ size appears to evolve largely independently through low pleiotropic mutations.

### Flower-size genes are under a mixed selection pattern in Arabidopsis thaliana

According to the most widely accepted theory about the evolution of selfing syndrome, the reduction of floral organ size results from the reallocation of resources no longer required for pollinator attraction toward other fitness-related traits. Under this hypothesis, new mutations reducing flower size should be favoured by natural selection and accumulate in selfing lineages. To test this, we estimated the ancestry (ancestral versus derived allele) and the total effect size of the alleles from all associated SNPs (see Methods). Contrary to expectations, we noted that more than three-quarters of petal area-derived alleles had a positive effect size, and this excess of positive effects among derived alleles was unique to petal area (**Figure 2a**). We further investigated whether it is a consequence of neutral processes or whether these alleles are under selection.

**Figure 2:**
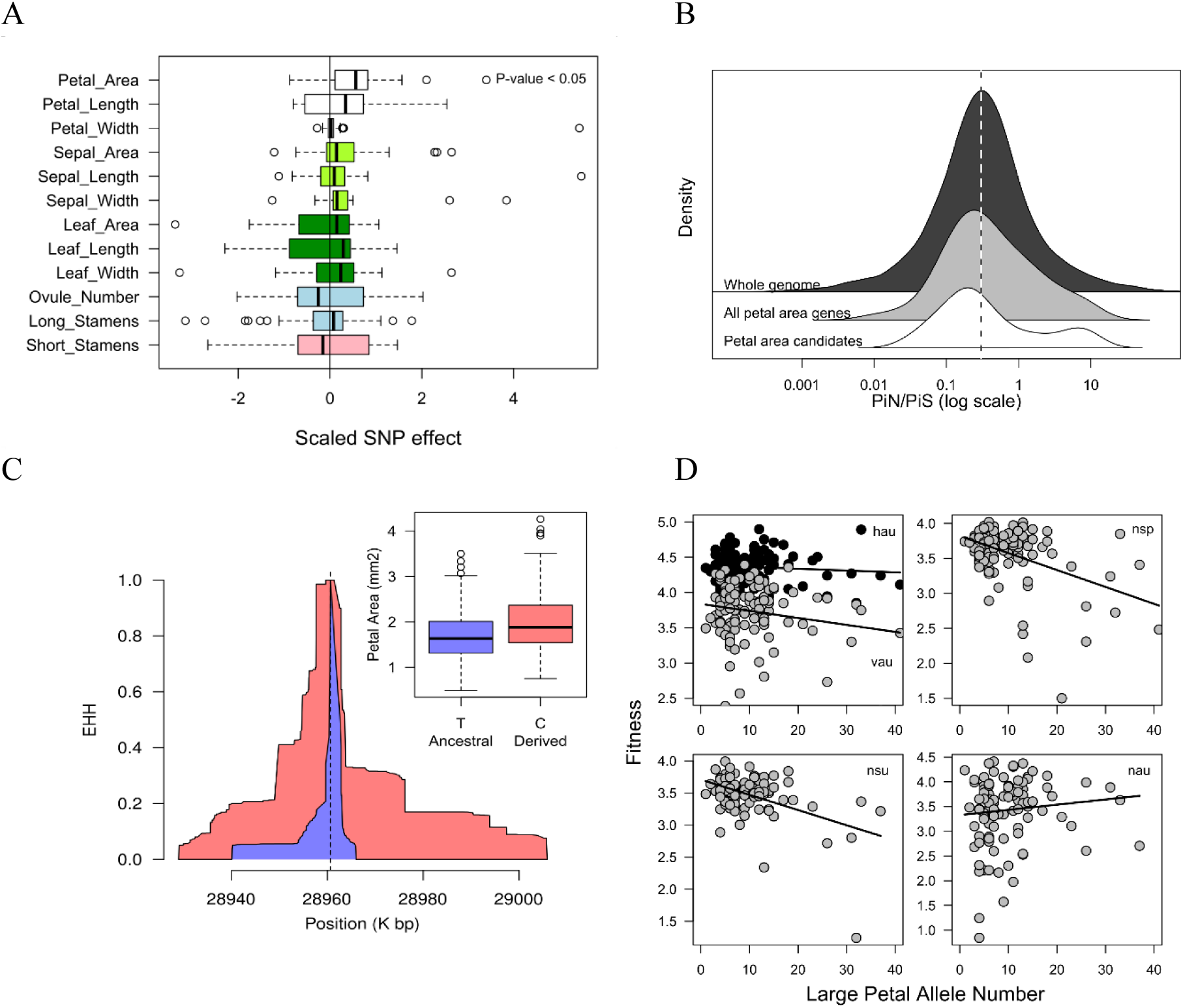
A variable flower size–seed production trade-off explains a mixed selection pattern on flower size in *A. thaliana*. **A)** Distribution of derived alleles’ total effect size from each associated SNPs identified in GWAS for each trait investigated. A p-value is given for trait distributions significantly different from 0 (vertical line). **B)** πN/πS distribution across all *Arabidopsis thaliana* genes, among the petal area genes and a priori candidate genes identified in the GWAS for petal area. The dashed line represents the whole genome average. **C)** Extended haplotype homozygosity around FLM lead SNP position. The red and blue integrals represent the degree of haplotype identity shared by individuals carrying the C (derived) and T (ancestral) alleles, respectively, as a function of distance to the lead SNP (dashed line). **D)** Relationship between the number of large petal allele and seed production across different experimental fields; hau, Halle autumn; vau, Valencia autumn; nsp, Norwich spring; nsu, Norwich summer; nau, and Norwich autumn. Black lines represent significant fits (anova) of a linear regressions.

To assess whether the fixation of derived alleles increasing flower size could result from genetic drift, we first computed genome-wide πN/πS, which contrasts the accumulation of non-synonymous (πN) and synonymous (πS) diversity to infer the strength and direction of selection. Genes containing SNPs associated with petal area exhibited a skewed πN/πS distribution that tended to fall below the genome-wide average (P-value < 0.1), indicating ongoing purifying selection rather than relaxed selection (**Figure 2b**). Other floral organ size traits showed no comparable pattern (**Figure S4**). While most petal candidate genes had πN/πS values below the genomic average, three genes, *AT1G36940*, *KIX9* and *FLM*, show elevated values within the top 5% of the genome-wide distribution, consistent with diversifying selection acting on their protein sequences.

We next examined whether derived alleles in petal area genes show evidence of positive selection. At a bi-allelic site, alleles under selection are expected to be associated with longer haplotypes due to incomplete selective sweeps. We estimated haplotype decay around each focal SNP using genome-wide extended haplotype homozygosity (EHH) (Klassmann & Gautier, 2022). The standardized ratio of derived to ancestral haplotype homozygosity (the integrated haplotype homozygosity score, iHS) identifies alleles with unusually long haplotypes for their frequency. Two petal area SNPs exhibited high |iHS| values, resulting from an extended haplotype homozygosity around the derived alleles, a sign of recent positive selection. The first, snp_1_10937139 (P = 0.013), increased petal area and likely affects the growth regulator JMJ18. The second, snp_5_870273 (P = 0.02), decreased petal area and likely influences AT5G03480, a poorly characterized RNA-binding protein expressed during early petal development. A third SNP, snp_1_28960616 (P = 0.07), showed a marginal signal of positive selection and likely affects FLM, a known flowering-time regulator that interacts with SVP to influence floral morphology (**Figure 2c**) (Posé et al., 2013). As a control, we repeated the analysis for all floral and leaf traits and quantified the enrichment of trait-associated SNPs within the top 0.05% of the genome-wide iHS distribution (Tsuchimatsu et al., 2020). Leaf size–associated SNPs showed a threefold enrichment, whereas no enrichment was detected for flower size (**Figure S5**).

Together, these results indicate a lack of consistent directional selection acting on the genetic determinants of flower size in *A. thaliana*. Neither purifying nor positive selection acts uniformly across petal size regulators, and no consistent directional trend is observable. The observed variation likely reflects spatially heterogeneous selection combined with neutral processes, accentuated by the species’ small effective population size and strong population structure. We next investigated whether phenotypic and genetic variation in floral organ size arises from spatially heterogeneous selective pressures across *A. thaliana*’s range.

### The influence of flower size on individual’s performance is environment dependent

The finding that alleles increasing flower size can be under positive selection in a selfing species challenges the resource allocation theory, which predicts higher fitness with reduced investment in floral organ growth in species such as *A. thaliana*. To investigate this apparent contradiction, we examined the relationship between petal size variation and plant fitness across populations and environments. We used fitness data from common garden experiments conducted at three sites, Valencia (Spain), Norwich (United Kingdom), and Halle (Germany) (Wilczek et al., 2014). Among the candidate SNPs associated with petal area, we counted the number of positive effect size alleles carried by each genotype, which provides a proxy for the genetic predisposition to produce larger petals. Consistent with expectations of the selfing syndrome, genotypes carrying more large-petal alleles generally performed worse than those carrying fewer large-petal alleles across most environments (**Figures 2d and S6**).

However, the relationship between the number of large petal alleles and fitness varied across sites and seasons. The relationship was negative in spring and summer in Norwich, but weakened in autumn in Halle and Valencia, and shift to a positive relationship in autumn in Norwich. Under more controlled experimental conditions, the negative relationship was significant in Madrid and Tuebingen in control treatments, whereas it weakened and became non-significant under drought treatments at both sites (**Figure S6**).

Overall, these results suggest that individual performance is affected by the genetic propensity to invest in petal growth, but the strength and form of this relationship depend on environmental conditions. Such context dependence likely generates heterogeneous selection on petal size across the geographic range of *A. thaliana*, possibly explaining the lack of unidirectional selection at the species level.

### Habitat suitability influences the selection regime at flower size loci

Our analyses so far suggest that the fitness costs and benefits of investing in floral growth in *A. thaliana* depend on local environmental conditions. To examine this further, we quantified habitat suitability at the collection sites of the studied populations. We used the *MaxEnt* software (Phillips et al., 2006) to model the species’ climatic niche based on bioclimatic variables (Karger et al., 2017), altitude (Danielson & Gesch, 2011), and a comprehensive occurrence dataset (Yim et al., 2022). From this model, we extracted the habitat suitability of each studied population (**Figures 3a and S7**) and the limiting environmental factors (**Figure S8**).

**Figure 3:**
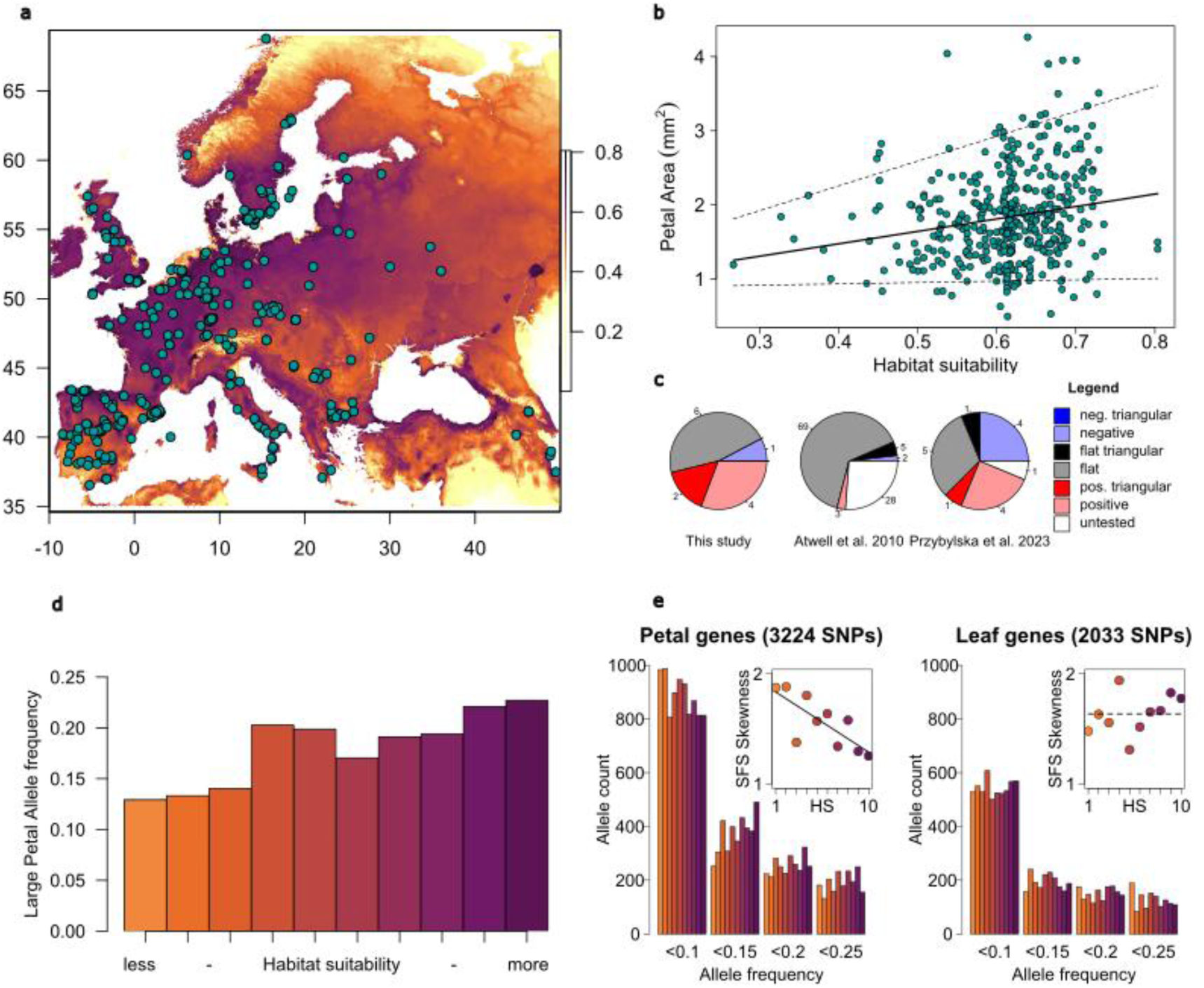
Habitat suitability shapes patterns of phenotype diversification across the geographical range of *A. thaliana*. **A)** Map of the studied European accessions. The orange to purple scale represents the modelled habitat suitability from less to more suitable, and the blue dots represent accessions’ collecting sites. **B)** Correlation between petal area and habitat suitability. Lines represent significant regressions: plain line, linear model; dashed lines, 0.05 and 0.95 quantile regressions. C) Relative proportion of different types of significant relationships between traits and habitat suitability in three independent datasets **D)** Large petal alleles’ frequency along habitat suitability clusters. **E)** Unfolded site frequency spectrum (SFS) among SNPs located in petal and leaf gene sets. The colours of the bars correspond to the habitat suitability classes. Inserts represent the relationship between the SFS’s shape and habitat suitability, where a plain line depicts significant and a dashed line insignificant correlation.

Across populations, petal area was positively associated with habitat suitability (**Figure 3b**). However, the relationship was triangular rather than linear: petal size variance increased toward more suitable environments, resembling the previously described relationship between flowering time and habitat suitability (Yim et al., 2022). Quantile regression confirmed this pattern, with the 0.05 and 0.95 quantile slopes differing significantly. This indicates that petal size evolution is more constrained at the climatic margins of the species’ range, where environmental stress limits expansion (Yim et al., 2022), while selection on flower size is more relaxed in favourable habitats.

To assess whether this triangular pattern is specific to petal area, we tested the relationship between habitat suitability and trait variance in multiple datasets (Atwell et al., 2010; Przybylska et al., 2023) (**Figure 3c**). None of the 107 traits in Atwell et al. (2010) showed significant heteroscedasticity, and only whole-plant dry mass in Przybylska et al. (2023) exhibited a similar positive triangular pattern. In our dataset, sepal length, in addition to petal area, displayed both a positive and triangular relationship with habitat suitability. Several other traits showed consistent but homoscedastic responses, including positive correlations for sepal size (this study), flowering time and leaf dry mass (Atwell et al., 2010; Przybylska et al., 2023), and negative correlations for leaf length, rosette diameter, fruit length, and leaf nitrogen content. Together, these results suggest that plant growth and life-history traits broadly align with habitat suitability, and that variance in floral organ size is particularly reduced under less favourable conditions.

To explore the evolutionary basis of this pattern, we examined how petal size alleles vary along the habitat suitability gradient. Consistent with the phenotypic trend, large-petal alleles increased in frequency in more suitable environments (**Figure 3d**). We tested whether this could reflect variation in purifying selection intensity across the species’ range. Purifying selection suppresses deleterious mutations, leading to an excess of rare derived alleles and a skewed site frequency spectrum (SFS). We computed the unfolded SFS for genes associated with petal and leaf traits to compare these effects. Low-frequency derived alleles (0.05 < frequency < 0.1) represented about one-fourth of SNPs in both gene sets (**Figure 3e**). The SFS of petal genes displayed significantly higher skewness in low-suitability habitats, indicating stronger purifying selection at the environmental margin. By contrast, SFS distributions for leaf genes did not differ along the gradient. These results indicate that the evolution of genes influencing flower size is more constrained in stressful environments, and purifying selection is relaxed in favourable habitats.

## Discussion

Contrary to predictions from resource allocation theories, our findings indicate that petal size variation in *Arabidopsis thaliana* is not governed by a uniform trade-off with fitness. Instead, it reflects a diversity of local selection pressures. Although producing large flowers can be costly, particularly in a species capable of generating up to a thousand flowers (Exposito-Alonso et al., 2018), this constraint is most pronounced at the margins of the species’ range where it favours small flowers. In contrast, in more favourable environments, the genetic propensity for larger petal size appears to be partially decoupled from fitness, allowing mutations that increase petal size to persist or even spread. Such spatial variation in selection likely contributes to the greater phenotypic variance in flower size among genotypes from climatically suitable regions.

The lack of genetic correlation and overlap in association signals suggests that petal size has largely evolved independently from other traits. However, our results do not completely exclude a role for life-history traits, such as flowering time, in petal size variation through pleiotropic effects. We detected a weak phenotypic correlation between flower size and flowering time, a trait known to be under positive selection in *A. thaliana* (Fournier-Level et al., 2022). Interestingly, among the few genes showing signatures of relaxed purifying selection and selective sweeps was *FLM*, a key regulator of flowering time that also influences plant growth (Hanemian et al., 2020). As *FLM* integrates temperature signals to control flowering, the absence of a clear GWAS signal at this locus does not rule out that selection on flowering time may indirectly affect flower size through the pleiotropic effects of shared regulatory pathways. Nevertheless, the highly polygenic nature of petal size, together with the identification of growth regulator candidates in genetic association, suggests that pleiotropy is unlikely to be a major determinant of flower size variation.

Although our study does not explicitly quantify gene-by-environment (G × E) interactions across the full environmental gradient occupied by *A. thaliana*, previous work indicates that G × E effects on flower size are modest, primarily affecting the magnitude rather than the direction of phenotypic responses (Wiszniewski et al., 2022). Consistent with this, petal size rankings among genotypes tend to remain relatively stable across environments, as reflected in the significant correlations between our predicted petal area and independent measurements of flower diameter at 17°C (R = 0.35, P-value < 0.01) and 23°C (R = 0.46, P-value < 0.01) (Wiszniewski et al., 2022). Moreover, the selective signatures observed at genes controlling flower size parallel the phenotypic trends, suggesting that functional constraints on small-flower alleles intensify toward the species’ climatic limits.

In addition to a relaxation of purifying selection, we detected evidence of positive selection acting on derived large-petal alleles, particularly in environmentally favourable regions. This suggests that an increase in petal size may confer adaptive advantages under certain conditions, or at least when environmental constraints on performance are minimal. The adaptive value of these alleles may relate to the persistence of a low but nonzero rate of cross-fertilisation in *A. thaliana* (Abbott & Gomes, 1989; Hoffmann et al., 2003), potentially favoured during post-glacial expansions northward, following secondary contact in the Iberian Peninsula (Fulgione & Hancock, 2018). Until today, the natural variation of selfing rate (s) across a large sample of *A. thaliana* populations has been reported once (Platt et al., 2010). Consistently, we detected a positive correlation between the outcrossing rate (log(1-s)) and petal area (R = 0.25, P-value = 0.12) and petal length (R = 0.32, P-value = 0.05) (**Figure S9**). While suggestive, these relationships should be interpreted cautiously due to the modest sample size (overlapping genotypes, N = 39) and the multiple-testing context, and will require validation through targeted analyses.

Other ecological factors may also promote larger flowers and should be explored in future studies, including historical interactions with floral enemies (Galen, 1999), protection against microbial and viral infections (Vannette, 2020), and the role of flower thermogenesis in pollen germination (van der Kooi et al., 2019). Pleiotropy and linked selection could further contribute to the emergence of large-flower alleles, especially as some petal size–associated variants lie within genes that influence other organs or developmental pathways (Hanemian et al., 2020). While our genetic correlation analyses indicate that such effects are limited, functional validation of causal mutations will be necessary to strictly disentangle developmental coupling from direct adaptive responses.

Overall, this study reveals striking geographic variation in investment in reproductive structures, particularly flower size, across the range of *A. thaliana*. Classic theory predicts that selfing lineages experience strong selection for reduced floral display, leading to the fixation of small-flower alleles due to their lower resource costs. Our results support this expectation only under environmentally stressful conditions, where resource limitation likely intensifies selection for smaller flowers. In contrast, in favourable environments, relaxed purifying selection reduces these constraints, enabling the persistence or even evolution of larger-flower variants. Together, these findings illustrate how habitat suitability and resource availability modulate the balance between selection and constraint, shaping the trajectory of phenotypic evolution in selfing species.

## Acknowledgments

We thank the Sicard, and Rosa groups for discussions and comments on the article. We thank Alexander Platt for valuable discussions and method sharing. The computation and data handling were provided by the Swedish National Infrastructure for Computing (SNIC) at Uppmax, partially funded by the Swedish Research Council through grant agreement no 2018-05973. This work was supported by the Swedish Research Council (grant 2018-04214) and the Novo Nordisk Foundation (NNF22OC0079830) to AS and JRL was supported by NIH award R35GM138300.

## Author Contributions

Conceptualization: KS, FV, CV and AS; investigations: KS, CFM, MJH, EG, PM and AE; Data analyses: KS and CFM; Materials and analysis tools contributions: DV and JRL; writing original draft KS and AS; writing –review & editing– All authors.

## References

Abbott, R. J., & Gomes, M. F. (1989). Population genetic structure and outcrossing rate of Arabidopsis thaliana (L.) Heynh. Heredity, 62(3), 411–418. 10.1038/hdy.1989.56

Abraham, M. C., Metheetrairut, C., & Irish, V. F. (2013). Natural Variation Identifies Multiple Loci Controlling Petal Shape and Size in Arabidopsis thaliana. PLOS ONE, 8(2), e56743. 10.1371/journal.pone.0056743

Alonso-Blanco, C., Andrade, J., Becker, C., Bemm, F., Bergelson, J., Borgwardt, K. M., Cao, J., Chae, E., Dezwaan, T. M., Ding, W., Ecker, J. R., Exposito-Alonso, M., Farlow, A., Fitz, J., Gan, X., Grimm, D. G., Hancock, A. M., Henz, S. R., Holm, S.,… Zhou, X. (2016). 1,135 Genomes Reveal the Global Pattern of Polymorphism in Arabidopsis thaliana. Cell, 166(2), 481–491. 10.1016/j.cell.2016.05.063

Atwell, S., Huang, Y. S., Vilhjálmsson, B. J., Willems, G., Horton, M., Li, Y., Meng, D., Platt, A., Tarone, A. M., Hu, T. T., Jiang, R., Muliyati, N. W., Zhang, X., Amer, M. A., Baxter, I., Brachi, B., Chory, J., Dean, C., Debieu, M.,… Nordborg, M. (2010). Genome-wide association study of 107 phenotypes in Arabidopsis thaliana inbred lines. Nature, 465(7298), 627–631. 10.1038/nature08800

Barrett, S. C. H., Arunkumar, R., & Wright, S. I. (2014). The demography and population genomics of evolutionary transitions to self-fertilization in plants. Philosophical Transactions of the Royal Society B: Biological Sciences, 369(1648), 20130344. 10.1098/rstb.2013.0344

Bechsgaard, J. S., Castric, V., Charlesworth, D., Vekemans, X., & Schierup, M. H. (2006). The Transition to Self-Compatibility in Arabidopsis thaliana and Evolution within S-Haplotypes over 10 Myr. Molecular Biology and Evolution, 23(9), 1741–1750. 10.1093/molbev/msl042

Burton, P. R., Clayton, D. G., Cardon, L. R., Craddock, N., Deloukas, P., Duncanson, A., Kwiatkowski, D. P., McCarthy, M. I., Ouwehand, W. H., Samani, N. J., Todd, J. A., Donnelly, P., Barrett, J. C., Burton, P. R., Davison, D., Donnelly, P., Easton, D., Evans, D., Leung, H.-T.,… Primary Investigators. (2007). Genome-wide association study of 14,000 cases of seven common diseases and 3,000 shared controls. Nature, 447(7145), Article 7145. 10.1038/nature05911

Buysse, S. F., Pérez, S. G., Puzey, J. R., Garrison, A., Bradburd, G. S., Oakley, C. G., Tonsor, S. J., Picó, F. X., Josephs, E. B., & Conner, J. K. (2024). Evaluating the Roles of Drift and Selection in Trait Loss along an Elevational Gradient (p. 2024.06.12.598645). bioRxiv. 10.1101/2024.06.12.598645

Caruso, C. M. (2006). Adaptive evolution: The ecological genetics of floral traits. Heredity, 97(2), 86–87. 10.1038/sj.hdy.6800853

Charlesworth, D., & Charlesworth, B. (1981). Allocation of resources to male and female functions in hermaphrodites. Biological Journal of the Linnean Society, 15(1), 57–74. 10.1111/j.1095-8312.1981.tb00748.x

Chen, Z., Boehnke, M., Wen, X., & Mukherjee, B. (2021). Revisiting the genome-wide significance threshold for common variant GWAS. G3 Genes|Genomes|Genetics, 11(2), jkaa056. 10.1093/g3journal/jkaa056

Cingolani, P. (2022). Variant Annotation and Functional Prediction: SnpEff. Methods in Molecular Biology (Clifton, N.J.), 2493, 289–314. 10.1007/978-1-0716-2293-3_19

Crane, P. R., Friis, E. M., & Pedersen, K. R. (1994). Palaeobotanical evidence on the early radiation of magnoliid angiosperms. In P. K. Endress & E. M. Friis (Eds.), Early Evolution of Flowers (pp. 51–72). Springer. 10.1007/978-3-7091-6910-0_4

Danielson, J. J., & Gesch, D. B. (2011). Global Multi-resolution Terrain Elevation Data 2010 (GMTED2010): U.S. Geo- logical Survey Open-File Report 2011 (p. 26). https://www.researchgate.net/publication/264159030_Global_Multi-resolution_Terrain_Elevation_Data_2010_GMTED2010

Dubos, C., Stracke, R., Grotewold, E., Weisshaar, B., Martin, C., & Lepiniec, L. (2010). MYB transcription factors in Arabidopsis. Trends in Plant Science, 15(10), 573–581. 10.1016/j.tplants.2010.06.005

Emms, D. M., & Kelly, S. (2015). OrthoFinder: Solving fundamental biases in whole genome comparisons dramatically improves orthogroup inference accuracy. Genome Biology, 16(1), 157. 10.1186/s13059-015-0721-2

Exposito-Alonso, M., Vasseur, F., Ding, W., Wang, G., Burbano, H. A., & Weigel, D. (2018). Genomic basis and evolutionary potential for extreme drought adaptation in Arabidopsis thaliana. Nature Ecology & Evolution, 2(2), 352–358. 10.1038/s41559-017-0423-0

Fournier-Level, A., Taylor, M. A., Paril, J. F., Martínez-Berdeja, A., Stitzer, M. C., Cooper, M. D., Roe, J. L., Wilczek, A. M., & Schmitt, J. (2022). Adaptive significance of flowering time variation across natural seasonal environments in Arabidopsis thaliana. New Phytologist, 234(2), 719–734. 10.1111/nph.17999

Friis, E. M., Pedersen, K., & Crane, P. (2006). Cretaceous angiosperm flowers: Innovation and evolution in plant reproduction. Palaeogeography, Palaeoclimatology, Palaeoecology, 232, 251–293. 10.1016/j.palaeo.2005.07.006

Friis, E. M., Pedersen, K. R., & Crane, P. R. (1994). Angiosperm floral structures from the Early Cretaceous of Portugal. In P. K. Endress & E. M. Friis (Eds.), Early Evolution of Flowers (pp. 31–49). Springer. 10.1007/978-3-7091-6910-0_3

Friis, E. M., Pedersen, K. R., & Crane, P. R. (1999). Early Angiosperm Diversification: The Diversity of Pollen Associated with Angiosperm Reproductive Structures in Early Cretaceous Floras from Portugal. Annals of the Missouri Botanical Garden, 86(2), 259. 10.2307/2666179

Fu, L., Wang, S., Liu, Z., Nijs, I., Ma, K., & Li, Z. (2010). Effects of resource availability on the trade-off between seed and vegetative reproduction. Journal of Plant Ecology, 3(4), 251–258. 10.1093/jpe/rtq017

Fulgione, A., & Hancock, A. M. (2018). Archaic lineages broaden our view on the history of Arabidopsis thaliana. New Phytologist, 219(4), 1194–1198. 10.1111/nph.15244

Galen, C. (1999). Why Do Flowers Vary?The functional ecology of variation in flower size and form within natural plant populations. BioScience, 49(8), 631–640. 10.2307/1313439

Gautier, M., & Vitalis, R. (2012). rehh: An R package to detect footprints of selection in genome-wide SNP data from haplotype structure. Bioinformatics, 28(8), 1176–1177. 10.1093/bioinformatics/bts115

Gómez-Ocampo, G., Crocco, C. D., Cascales, J., Oklestkova, J., Tarkowská, D., Strnad, M., Mora-Garcia, S., Pruneda-Paz, J. L., Blazquez, M. A., & Botto, J. F. (2023). BBX21 Integrates Brassinosteroid Biosynthesis and Signaling in the Inhibition of Hypocotyl Growth under Shade. Plant and Cell Physiology, pcad126. 10.1093/pcp/pcad126

Gonzalez, N., Pauwels, L., Baekelandt, A., De Milde, L., Van Leene, J., Besbrugge, N., Heyndrickx, K. S., Cuéllar Pérez, A., Durand, A. N., De Clercq, R., Van De Slijke, E., Vanden Bossche, R., Eeckhout, D., Gevaert, K., Vandepoele, K., De Jaeger, G., Goossens, A., & Inzé, D. (2015). A Repressor Protein Complex Regulates Leaf Growth in Arabidopsis. The Plant Cell, 27(8), 2273–2287. 10.1105/tpc.15.00006

Hanemian, M., Vasseur, F., Marchadier, E., Gilbault, E., Bresson, J., Gy, I., Violle, C., & Loudet, O. (2020). Natural variation at FLM splicing has pleiotropic effects modulating ecological strategies in Arabidopsis thaliana. Nature Communications, 11(1), 4140. 10.1038/s41467-020-17896-w

Hoffmann, M. H., Bremer, M., Schneider, K., Burger, F., Stolle, E., & Moritz, G. (2003). Flower Visitors in a Natural Population of Arabidopsis thaliana. Plant Biology, 5(5), 491–494. 10.1055/s-2003-44784

Huxley, J. S., Churchill, F. B., & Strauss, R. E. (1993). Problems of Relative Growth. Johns Hopkins University Press. 10.56021/9780801846595

Johnson, K., & Lenhard, M. (2011). Genetic control of plant organ growth. New Phytologist, 191(2), 319–333. 10.1111/j.1469-8137.2011.03737.x

Karger, D. N., Conrad, O., Böhner, J., Kawohl, T., Kreft, H., Soria-Auza, R. W., Zimmermann, N. E., Linder, H. P., & Kessler, M. (2017). Climatologies at high resolution for the earth’s land surface areas. Scientific Data, 4, 170122. 10.1038/sdata.2017.122

Katoh, K., Misawa, K., Kuma, K., & Miyata, T. (2002). MAFFT: A novel method for rapid multiple sequence alignment based on fast Fourier transform. Nucleic Acids Research, 30(14), 3059. 10.1093/nar/gkf436

Keightley, P. D., & Jackson, B. C. (2018). Inferring the Probability of the Derived vs. The Ancestral Allelic State at a Polymorphic Site. Genetics, 209(3), 897–906. 10.1534/genetics.118.301120

Klassmann, A., & Gautier, M. (2022). Detecting selection using extended haplotype homozygosity (EHH)-based statistics in unphased or unpolarized data. PLoS ONE, 17(1). 10.1371/JOURNAL.PONE.0262024

Lee, D. J., Kim, S., Ha, Y.-M., & Kim, J. (2008). Phosphorylation of Arabidopsis response regulator 7 (ARR7) at the putative phospho-accepting site is required for ARR7 to act as a negative regulator of cytokinin signaling. Planta, 227(3), 577–587. 10.1007/s00425-007-0640-x

Li, X., Zhang, Y., Yang, S., Wu, C., Shao, Q., & Feng, X. (2020). The genetic control of leaf and petal allometric variations in Arabidopsis thaliana. BMC Plant Biology, 20(1), 1–11. 10.1186/s12870-020-02758-w

Lopes, A. L., Moreira, D., Pereira, A. M., Ferraz, R., Mendes, S., Pereira, L. G., Colombo, L., & Coimbra, S. (2023). AGPs as molecular determinants of reproductive development. Annals of Botany, 131(5), 827–838. 10.1093/aob/mcad046

MacTavish, R., & Anderson, J. T. (2020). Resource availability alters fitness trade-offs: Implications for evolution in stressful environments. American Journal of Botany, 107(2), 308–318. 10.1002/ajb2.1417

Maki, Y., Soejima, H., Sugiyama, T., Watahiki, M. K., Sato, T., & Yamaguchi, J. (2022). 3-Phenyllactic acid is converted to phenylacetic acid and induces auxin-responsive root growth in Arabidopsis plants. Plant Biotechnology, 39(2), 111–117. 10.5511/plantbiotechnology.21.1216a

Omidbakhshfard, M. A., Proost, S., Fujikura, U., & Mueller-Roeber, B. (2015). Growth-Regulating Factors (GRFs): A Small Transcription Factor Family with Important Functions in Plant Biology. Molecular Plant, 8(7), 998–1010. 10.1016/j.molp.2015.01.013

Peng, R. K., Victor Chernozhukov, Xuming He, Limin (Ed.). (2017). Handbook of Quantile Regression. Chapman and Hall/CRC. 10.1201/9781315120256

Pérez-Harguindeguy, N., Díaz, S., Garnier, E., Lavorel, S., Poorter, H., Jaureguiberry, P., Bret-Harte, M. S., Cornwell, W. K., Craine, J. M., Gurvich, D. E., Urcelay, C., Veneklaas, E. J., Reich, P. B., Poorter, L., Wright, I. J., Ray, P., Enrico, L., Pausas, J. G., de Vos, A. C.,… Cornelissen, J. H. C. (2013). New handbook for standardised measurement of plant functional traits worldwide. Australian Journal of Botany, 61(3), 167. 10.1071/BT12225

Phillips, S. J., Anderson, R. P., & Schapire, R. E. (2006). Maximum entropy modeling of species geographic distributions. Ecological Modelling, 190(3), 231–259. 10.1016/j.ecolmodel.2005.03.026

Platt, A., Horton, M., Huang, Y. S., Li, Y., Anastasio, A. E., Mulyati, N. W., Ågren, J., Bossdorf, O., Byers, D., Donohue, K., Dunning, M., Holub, E. B., Hudson, A., Le Corre, V., Loudet, O., Roux, F., Warthmann, N., Weigel, D., Rivero, L.,… Borevitz, J. O. (2010). The Scale of Population Structure in Arabidopsis thaliana. PLoS Genetics, 6(2), e1000843. 10.1371/journal.pgen.1000843

Przybylska, M. S., Violle, C., Vile, D., Scheepens, J. F., Lacombe, B., Le Roux, X., Perrier, L., Sales-Mabily, L., Laumond, M., Vinyeta, M., Moulin, P., Beurier, G., Rouan, L., Cornet, D., & Vasseur, F. (2023). AraDiv: A dataset of functional traits and leaf hyperspectral reflectance of Arabidopsis thaliana. Scientific Data, 10(1), Article 1. 10.1038/s41597-023-02189-w

Purcell, S., Neale, B., Todd-Brown, K., Thomas, L., Ferreira, M. A. R., Bender, D., Maller, J., Sklar, P., de Bakker, P. I. W., Daly, M. J., & Sham, P. C. (2007). PLINK: A Tool Set for Whole-Genome Association and Population-Based Linkage Analyses. The American Journal of Human Genetics, 81(3), 559–575. 10.1086/519795

Roddy, A. B., Théroux-Rancourt, G., Abbo, T., Benedetti, J. W., Brodersen, C. R., Castro, M., Castro, S., Gilbride, A. B., Jensen, B., Jiang, G.-F., Perkins, J. A., Perkins, S. D., Loureiro, J., Syed, Z., Thompson, R. A., Kuebbing, S. E., & Simonin, K. A. (2020). The Scaling of Genome Size and Cell Size Limits Maximum Rates of Photosynthesis with Implications for Ecological Strategies. International Journal of Plant Sciences, 181(1), 75–87. 10.1086/706186

Sabeti, P. C., Reich, D. E., Higgins, J. M., Levine, H. Z. P., Richter, D. J., Schaffner, S. F., Gabriel, S. B., Platko, J. V., Patterson, N. J., McDonald, G. J., Ackerman, H. C., Campbell, S. J., Altshuler, D., Cooper, R., Kwiatkowski, D., Ward, R., & Lander, E. S. (2002). Detecting recent positive selection in the human genome from haplotype structure. Nature, 419(6909), 832–837. 10.1038/nature01140

Sasaki, E., Frommlet, F., & Nordborg, M. (2018). GWAS with Heterogeneous Data: Estimating the Fraction of Phenotypic Variation Mediated by Gene Expression Data. G3 Genes|Genomes|Genetics, 8(9), 3059–3068. 10.1534/g3.118.200571

Sauret-Güeto, S., Schiessl, K., Bangham, A., Sablowski, R., & Coen, E. (2013). JAGGED Controls Arabidopsis Petal Growth and Shape by Interacting with a Divergent Polarity Field. PLoS Biology, 11(4), e1001550. 10.1371/journal.pbio.1001550

Schindelin, J., Arganda-Carreras, I., Frise, E., Kaynig, V., Longair, M., Pietzsch, T., Preibisch, S., Rueden, C., Saalfeld, S., Schmid, B., Tinevez, J.-Y., White, D. J., Hartenstein, V., Eliceiri, K., Tomancak, P., & Cardona, A. (2012). Fiji: An open-source platform for biological-image analysis. Nature Methods, 9(7), Article 7. 10.1038/nmeth.2019

Shimizu, K. K., & Tsuchimatsu, T. (2015). Evolution of Selfing: Recurrent Patterns in Molecular Adaptation. Annual Review of Ecology, Evolution, and Systematics, 46(1), 593–622. 10.1146/annurev-ecolsys-112414-054249

Sicard, A., & Lenhard, M. (2011). The selfing syndrome: A model for studying the genetic and evolutionary basis of morphological adaptation in plants. Annals of Botany, 107(9), 1433–1443. 10.1093/aob/mcr023

Sicard, A., Stacey, N., Hermann, K., Dessoly, J., Neuffer, B., Bäurle, I., & Lenhard, M. (2011). Genetics, evolution, and adaptive significance of the selfing syndrome in the genus Capsella. The Plant Cell, 23(9), 3156–3171. 10.1105/tpc.111.088237

Slotte, T. (2014). The impact of linked selection on plant genomic variation. Briefings in Functional Genomics, 13(4), 268–275. 10.1093/bfgp/elu009

Smith, J. M., & Haigh, J. (1974). The hitch-hiking effect of a favourable gene. Genetics Research, 23(1), 23–35. 10.1017/S0016672300014634

Stebbins, G. L. (1957). Self Fertilization and Population Variability in the Higher Plants. The American Naturalist, 91(861), 337–354. 10.1086/281999

Stephan, W. (2019). Selective Sweeps. Genetics, 211(1), 5–13. 10.1534/genetics.118.301319

Tang, C., Toomajian, C., Sherman-Broyles, S., Plagnol, V., Guo, Y.-L., Hu, T. T., Clark, R. M., Nasrallah, J. B., Weigel, D., & Nordborg, M. (2007). The evolution of selfing in Arabidopsis thaliana. Science (New York, N.Y.), 317(5841), 1070–1072. 10.1126/science.1143153

Troth, A., Puzey, J. R., Kim, R. S., Willis, J. H., & Kelly, J. K. (2018). Selective trade-offs maintain alleles underpinning complex trait variation in plants. Science, 361(6401), 475–478. 10.1126/science.aat5760

Tsuchimatsu, T., Kakui, H., Yamazaki, M., Marona, C., Tsutsui, H., Hedhly, A., Meng, D., Sato, Y., Städler, T., Grossniklaus, U., Kanaoka, M. M., Lenhard, M., Nordborg, M., & Shimizu, K. K. (2020). Adaptive reduction of male gamete number in the selfing plant Arabidopsis thaliana. Nature Communications 2020 11:1, 11(1), 1–9. 10.1038/s41467-020-16679-7

Tuller, J., Marquis, R. J., Andrade, S. M. M., Monteiro, A. B., & Faria, L. D. B. (2018). Trade-offs between growth, reproduction and defense in response to resource availability manipulations. PLOS ONE, 13(8), e0201873. 10.1371/journal.pone.0201873

Vannette, R. L. (2020). The Floral Microbiome: Plant, Pollinator, and Microbial Perspectives. Annual Review of Ecology, Evolution, and Systematics, 51(1), 363–386. 10.1146/annurev-ecolsys-011720-013401

Vlieghe, K., Boudolf, V., Beemster, G. T. S., Maes, S., Magyar, Z., Atanassova, A., Engler, J. de A., Groodt, R. D., Inzé, D., & Veylder, L. D. (2005). The DP-E2F-like Gene DEL1 Controls the Endocycle in Arabidopsis thaliana. Current Biology, 15(1), 59–63. 10.1016/j.cub.2004.12.038

Voight, B. F., Kudaravalli, S., Wen, X., & Pritchard, J. K. (2006). A Map of Recent Positive Selection in the Human Genome. PLOS Biology, 4(3), e72. 10.1371/journal.pbio.0040072

Warton, D. I., Wright, I. J., Falster, D. S., & Westoby, M. (2006). Bivariate line-fitting methods for allometry. Biological Reviews, 81(02), 259. 10.1017/S1464793106007007

Wilczek, A. M., Cooper, M. D., Korves, T. M., & Schmitt, J. (2014). Lagging adaptation to warming climate in Arabidopsis thaliana. Proceedings of the National Academy of Sciences, 111(22), 7906–7913. 10.1073/pnas.1406314111

Wiszniewski, A., Uberegui, E., Messer, M., Sultanova, G., Borghi, M., Duarte, G. T., Vicente, R., Sageman-Furnas, K., Fernie, A. R., Nikoloski, Z., & Laitinen, R. A. E. (2022). Temperature-mediated flower size plasticity in Arabidopsis. iScience, 25(11), 105411. 10.1016/J.ISCI.2022.105411

Worley, A. C., & Barrett, S. C. H. (2000). Evolution of Floral Display in Eichhornia Paniculata (pontederiaceae): Direct and Correlated Responses to Selection on Flower Size and Number. Evolution, 54(5), 1533–1545. 10.1111/j.0014-3820.2000.tb00699.x

Woźniak, N. J., Kappel, C., Marona, C., Altschmied, L., Neuffer, B., & Sicard, A. (2020). A Similar Genetic Architecture Underlies the Convergent Evolution of the Selfing Syndrome in *Capsella*. The Plant Cell, 32(4), 935–949. 10.1105/tpc.19.00551

Woźniak, N. J., & Sicard, A. (2018). Evolvability of flower geometry: Convergence in pollinator-driven morphological evolution of flowers. Seminars in Cell & Developmental Biology, Shape and Form in Plant Development, 79, 3–15. 10.1016/j.semcdb.2017.09.028

Yim, C., Bellis, E. S., DeLeo, V. L., Gamba, D., Muscarella, R., & Lasky, J. R. (2022). Climate biogeography of Arabidopsis thaliana: Linking distribution models, individual performance, and life history (p. 2022.03.06.483202). bioRxiv. 10.1101/2022.03.06.483202

Zhou, X., Carbonetto, P., & Stephens, M. (2013). Polygenic modeling with bayesian sparse linear mixed models. PLoS Genetics, 9(2), e1003264. 10.1371/journal.pgen.1003264

Zhou, X., & Stephens, M. (2012). Genome-wide efficient mixed-model analysis for association studies. Nature Genetics, 44(7), 821–824.

